# Fragment anchoring for metabolite annotation beyond whole-spectrum similarity

**DOI:** 10.64898/2026.11.11.766011

**Authors:** Young Jae Park, Ji Hyun Moon, Ki Tae Suk, Do Yup Lee

## Abstract

Many untargeted metabolomics features remain unannotated, and chemically related metabolites can have dissimilar tandem mass spectra. We present ANCHor, a framework for interpreting features of interest using shared fragment evidence. ANCHor selects fragments of a feature that recur among similar atlas spectra but are rare in background spectra, then evaluates matching experimental features using mass differences and transformation rules. Using 1,021 authentic standards as features of interest, ANCHor generated at least one anchor for 76.9%. We tested whether L-methionine anchors retain related spectra despite low overall similarity. Of 166 structurally related spectra, ANCHor retained 147, including 93 with modified cosine similarity <0.70 to L-methionine. In mycotoxin-treated hepatic spheroids, ANCHor prioritized putative transformation products, including low-similarity candidates. For library-unmatched fecal features in alcohol-associated liver disease, ANCHor found annotated references that guided identification of N¹,N¹²-diacetylspermine, confirmed with an authentic standard, and supported a tryptophan–pyruvate product hypothesis with a reaction-generated reference. ANCHor thus supports testable structural hypotheses for library-unmatched features within atlas coverage.

## Introduction

LC–MS/MS-based untargeted metabolomics is widely used to profile diverse small-molecule signals in biological samples and to explore chemical changes associated with biological states or external exposures^1–3^. However, although a single experiment can detect thousands to tens of thousands of features, many detected signals remain chemically unassigned. For features of interest, identifying plausible links to known metabolites or chemical classes remains a major bottleneck^3–6^.

Spectral library matching provides relatively high-confidence annotation by directly comparing experimental MS/MS spectra with reference or authentic standard spectra^7^. However, its application is limited to compounds for which reference spectra are available. As a result, many features in untargeted metabolomics datasets remain unmatched or are supported only by partial structural information^7–10^. This limitation has motivated approaches that interpret unknown features in relation to annotated features, rather than annotating each feature in isolation, thereby narrowing the set of plausible annotations^11,12^.

Molecular networking has been widely used as a representative approach for structural interpretation of large-scale LC–MS/MS data by connecting features based on MS/MS spectral similarity and grouping them into molecular families^13–16^. This approach is particularly useful for visually organizing features with similar fragmentation patterns and propagating known annotations to neighboring features^17,18^. However, because feature-to-feature connections in molecular networking are largely determined by global spectral similarity metrics, such as cosine or modified cosine similarity, metabolites linked by known biochemical transformations may not always form direct network edges^19^. For example, biotransformations such as conjugation, oxidation/reduction, or methylation can alter both precursor mass and fragmentation behavior, causing related metabolites to show low spectral similarity and become separated in the network^19–21^. In addition, missing intermediate nodes or low spectral coverage can leave some features as singletons or disconnected subnetworks^16,20^. Therefore, a complementary approach is needed to identify and interpret features related to a feature of interest and to explicitly summarize the evidence supporting those relationships.

Here, we present ANCHor (Atlas-iNformed CHemical anchoring), a seed-centric metabolite interpretation pipeline designed to support unknown feature annotation. In ANCHor, a user-specified feature of interest is referred to as the seed. ANCHor first retrieves seed-neighbor spectra from a large-scale spectral atlas and selects seed anchors, defined as seed-associated MS/MS fragment signals that are present in the seed spectrum, recurrent among seed-neighbor spectra, and rare in background spectra. These seed anchors are then projected onto the user’s local feature list to identify projection-retained features, and seed–feature relationships are interpreted using knowledge-based transformation rules. Rather than relying solely on pairwise spectral similarity, ANCHor combines shared-anchor evidence with mass-difference and transformation-rule evidence to generate testable annotation hypotheses.

The primary goal of ANCHor is not simply to add feature-to-feature links, but to generate testable annotation hypotheses for library-unmatched features. This is particularly relevant when statistically significant features lack library matches. For such features, ANCHor identifies plausible relationships to annotated metabolites and evaluates whether those relationships are supported by shared-anchor and mass-difference evidence. By organizing this evidence around the seed feature, ANCHor can help prioritize unknown features and guide validation of putative metabolite identities or biotransformation relationships.

ANCHor was benchmarked using an in-house reference library of authentic standards generated with the same analytical workflow used for the experimental datasets. We then applied ANCHor to a mycotoxin-treated HepG2 spheroid model to examine whether the workflow could trace the biotransformation of an external compound in a cell-based hepatic metabolism model. Finally, we applied ANCHor to a human fecal metabolomics cohort comparing alcohol-associated liver disease (ALD) and control groups. In this cohort application, a statistically significant library-unmatched feature was assigned an annotation hypothesis using ANCHor-derived chemical evidence and was subsequently confirmed by comparison with a synthesized reference compound based on matching retention time and MS/MS spectra. Together, these analyses demonstrate that ANCHor provides a general framework for seed-centered feature interpretation, spanning anchor discovery in reference spectra, prioritization of biotransformation-related features, and generation of experimentally testable annotation hypotheses for library-unmatched features.

## Results

### Overview of the ANCHor workflow

ANCHor was designed as a three-step workflow that begins with a user-specified feature of interest, hereafter referred to as the seed (Fig. 1). The workflow consists of anchoring, projection, and interpretation. In the anchoring step, ANCHor uses DreaMS similarity to retrieve seed-neighbor spectra from a precursor-m/z-matched subset of a large-scale spectral atlas. It then selects seed anchors from the seed spectrum. These anchors are MS/MS fragment signals that recur among seed-neighbor spectra but remain rare in background spectra (Fig. 1a). This step defines atlas-supported fragment evidence for the seed using a precursor-m/z-matched retrieval context. This comparison allows ANCHor to identify seed fragments that recur preferentially in DreaMS-similar spectra relative to mass-matched dissimilar spectra.

**Fig. 1.**
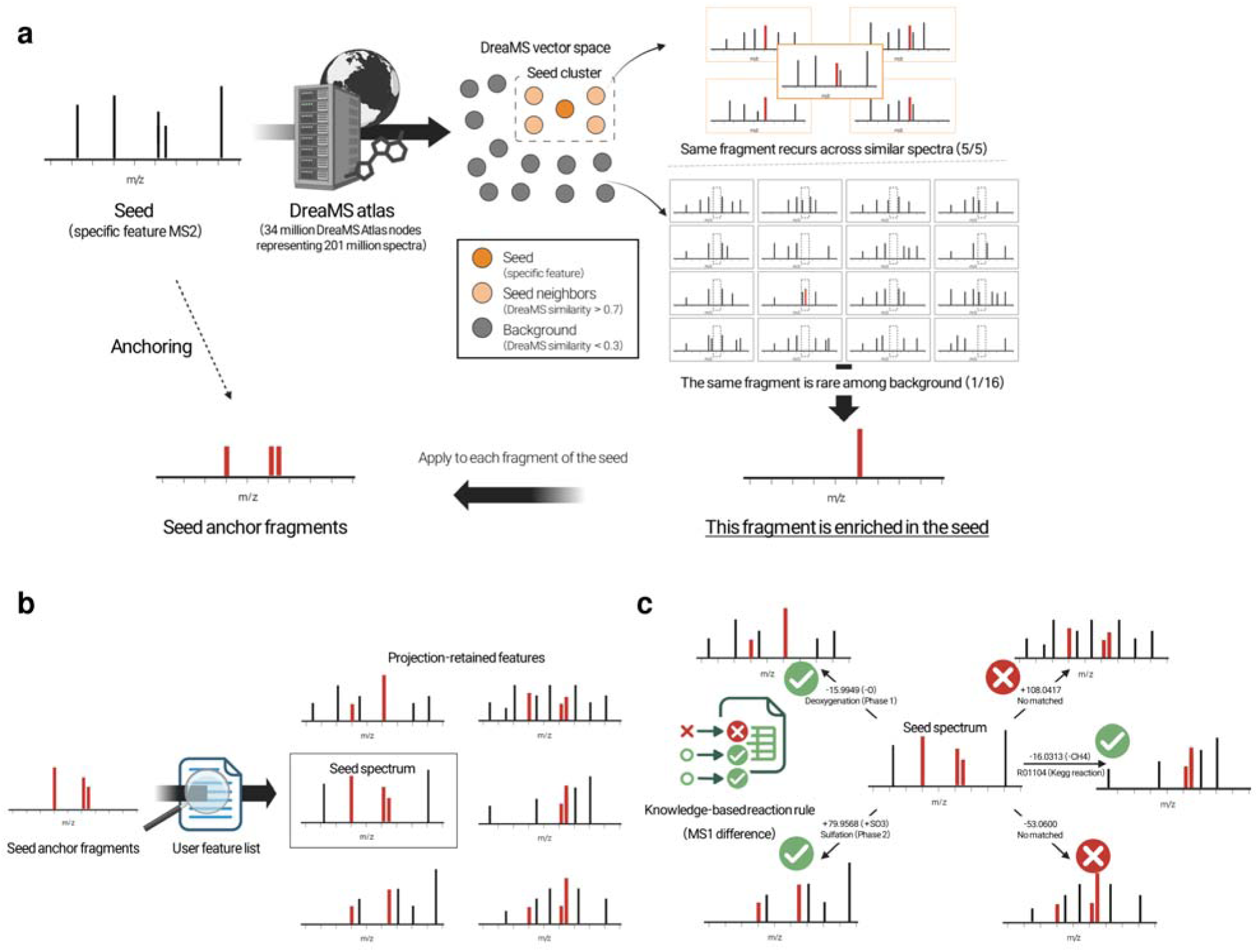
Overview of ANCHor for seed-centered metabolite interpretation. **a** Anchoring. ANCHor (AtlasiNformed CHemical anchoring) retrieved DreaMS atlas spectra with embedding similarity ≥0.70 to the seed as seed-neighbor spectra. Seed-associated MS/MS fragments recurrent among these spectra but rare in background spectra were selected as anchors. The experimental seed was excluded from atlas occurrence counts. **b** Projection. Selected seed anchors were matched to MS/MS spectra in the local feature list. Features meeting the shared-anchor and coverage criteria were retained as projection-retained features. **c** Interpretation. Projection-retained features were evaluated using knowledge-based transformation rules, mass differences and shared-anchor evidence.

In the projection step, the selected seed anchors are projected onto all MS/MS spectra in the user’s dataset. In contrast to atlas retrieval, projection is not restricted to precursor-m/z-matched spectra. This broader search is based on the premise that chemically related molecules may retain a subset of seed-associated fragments even when their precursor m/z values and overall MS/MS spectra differ. Features meeting the predefined anchor-sharing criteria are retained for interpretation and termed projection-retained features (Fig. 1b).

Shared-anchor evidence alone does not specify how a projection-retained feature may be chemically related to the seed. In the interpretation step, ANCHor therefore evaluates the neutral mass difference between each projection-retained feature and the seed against knowledge-based transformation rules (Fig. 1c).

Overall, ANCHor takes a feature table, an MGF file, and a user-specified seed feature ID as input. It returns projection-retained features with shared-anchor evidence, mass-difference evidence, and transformation-rule-based annotation hypotheses.

### Evaluation of anchor discovery and projection

To evaluate whether ANCHor could generate seed anchors across diverse chemical categories, we applied the anchoring step to an in-house authentic standard library acquired in positive ionization mode. The MS/MS spectrum of each of 1,021 unique standards was used as the seed. ANCHor generated at least one final seed anchor for 785 standards (76.9%), with a median of 4 final anchors among these seeds.

Anchor-generation rates were then summarized at the superclass, class, and subclass levels based on ClassyFire classifications^22^ (Fig. 2a). Among the five most abundant categories at each classification level, rates ranged from 74.3% to 81.7% for superclasses, 73.8% to 94.3% for classes, and 65.7% to 100% for subclasses. For the remaining categories, anchor-generation rates were 73.2% at the class level and 75.2% at the subclass level. Several lipid-related categories showed high anchor-generation rates, including lipids and lipid-like molecules (81.7%), fatty acyls (81.9%), and steroids and steroid derivatives (94.3%), although fatty acids and conjugates showed a lower rate of 65.7%.

**Fig. 2.**
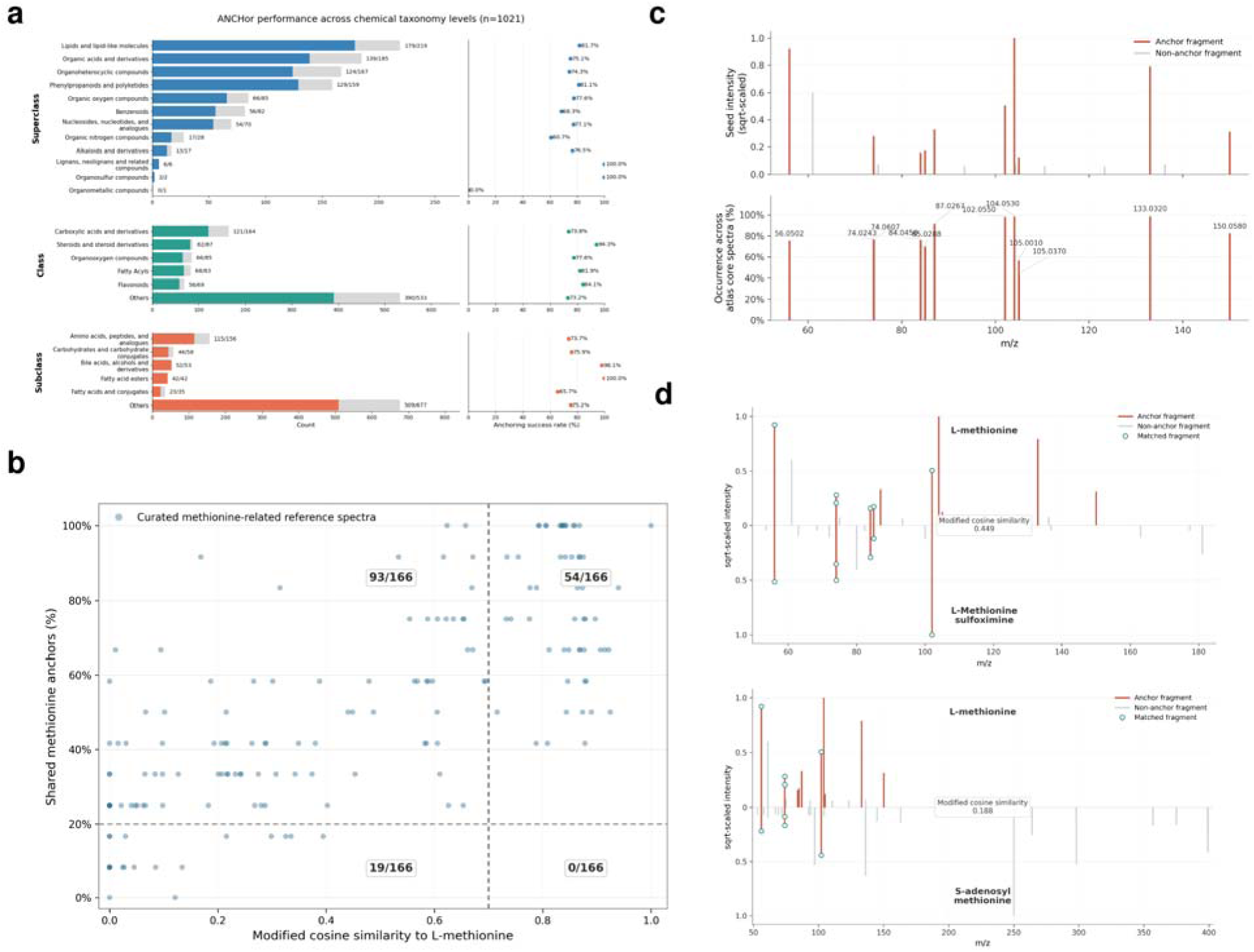
Evaluation of anchor discovery and seed-anchor projection. **a** Anchoring success across superclass, class and subclass levels in an in-house library of 1,021 authentic standards. **b** Comparison of modified cosine similarity to L-methionine and methionine-anchor sharing across 166 curated methionine-related reference MS/MS spectra. Dashed lines indicate the modified cosine threshold (≥0.70) and seed-anchor-sharing threshold (≥20%). **C** Experimental L-methionine MS/MS spectrum and atlas-derived pseudo-spectrum. Pseudo-spectrum peak heights indicate fragment occurrence among methionine seed-neighbor spectra. **d** Mirror plots comparing L-methionine with representative methionine-related reference spectra retained by seed-anchor sharing.

Anchors were not generated for 236 standards (23.1%). For 142 standards, no atlas spectra meeting the DreaMS similarity threshold of 0.70 were retrieved within the precursor-m/z window. For the remaining 94 standards, seed-neighbor spectra were available, but no fragment passed the final anchor-selection criteria. These failures distinguish the absence of qualifying atlas neighbors from failure to select an anchor despite their availability. Nevertheless, anchors were generated for the majority of authentic standards, indicating that ANCHor can derive seed-associated fragment evidence across diverse chemical categories represented in the library.

We next used L-methionine as a representative seed and assembled a structurally defined panel of 166 annotated MS/MS spectra from reference spectral libraries (GNPS, MoNA and NIST)^14,23,24^. Panel entries were selected by matching their annotated SMILES against a manually curated SMARTS pattern encoding the methionine carbon–sulfur substructure. This panel was used to evaluate whether methionine-derived anchors could retain structurally related spectra of compounds sharing the target substructure. Anchoring of L-methionine yielded 12 final seed anchors (Supplementary Fig. 1). Of the 166 annotated spectra in the panel, 54 (33%) had a modified cosine similarity ≥ 0.70 to L-methionine, a threshold commonly used in molecular networking^15,19^. In contrast, 147 spectra (89%) met the predefined ANCHor projection criterion (Fig. 2b). Notably, the ANCHor projection criterion retained a substantially larger fraction of the structurally defined panel, including many spectra with low modified cosine similarity to L-methionine, indicating that seed-anchor evidence provides information complementary to whole-spectrum similarity. The full ANCHor projection criterion was predefined and applied throughout the study rather than optimized using the methionine panel. Its minimum shared-anchor requirement reflects the premise that structurally related metabolites may preserve only part of the seed-derived fragment pattern rather than the complete anchor set.

Methionine-derived anchors were visualized by comparing the L-methionine spectrum with an atlas-derived pseudo-spectrum. In the pseudo-spectrum, peak height represents how frequently each fragment occurred among methionine seed-neighbor spectra (Fig. 2c). A relatively intense seed peak at m/z 61.0106 occurred in 85.95% of seed-neighbor spectra but also in 29.32% of BG1 spectra, exceeding the <15% background-occurrence threshold, and was therefore excluded. This example illustrates that anchor selection requires both recurrence among seed-neighbor spectra and rarity in background spectra, rather than high intensity in the seed spectrum alone. Representative mirror plots illustrate how individual methionine-related spectra shared multiple seed anchors despite low modified cosine similarity. L-methionine sulfoximine shared 6 of 12 methionine anchors with a modified cosine similarity of 0.449 to L-methionine (Fig. 2d). S-adenosyl-L-methionine, which is metabolically adjacent to L-methionine, shared 4 of 12 anchors (33%) despite a modified cosine similarity of 0.188 (Supplementary Fig. 2). Neither spectrum reached the modified cosine threshold of 0.70 for a direct network edge, but both met the ANCHor projection criteria.

Overall, the authentic-standard benchmark showed that seed anchors could be generated across diverse chemical categories, and the methionine panel showed that the ANCHor projection criterion could retain structurally related spectra even when pairwise modified cosine similarity was low.

### Mycotoxin biotransformation in hepatic spheroids

We next applied ANCHor to a zearalenone (ZEN)-treated HepG2 spheroid model to evaluate whether the workflow could prioritize features associated with xenobiotic biotransformation. ZEN was selected as a model mycotoxin because its reported metabolism involves multiple reaction types, including reduction and conjugation^25,26^. HepG2 spheroids were used to provide a more metabolically competent hepatic context than conventional 2D monolayer culture^27,28^. Conditioned medium from ZEN-treated HepG2 spheroids was analyzed by untargeted LC–MS/MS, yielding 2,343 detected features, and the ZEN feature was used as the seed for ANCHor analysis (Fig. 3a).

**Fig. 3.**
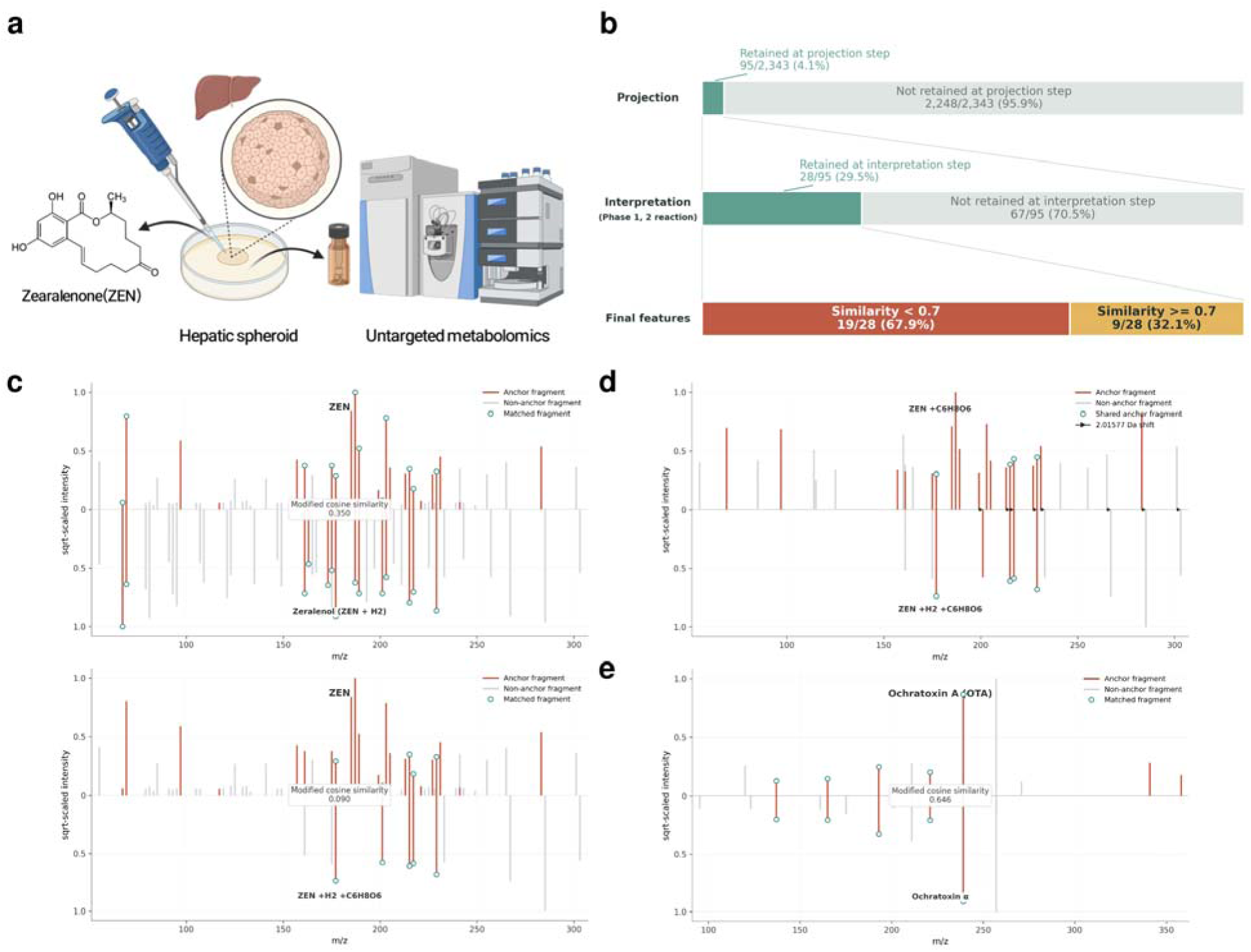
ANCHor-assisted annotation of mycotoxin biotransformation. **a** Experimental workflow for the zearalenone (ZEN) model. HepG2 spheroids were exposed to ZEN, and conditioned medium was analyzed by untargeted LC–MS/MS. ZEN was used as the seed for ANCHor analysis. **b** Stepwise prioritization of ZEN-related features. **c** Mirror plots comparing ZEN with putative zearalenol (upper comparison; modified cosine similarity, 0.350) and putative zearalenol glucuronide (lower comparison; 0.090). **d** Comparison of putative zearalenone glucuronide and putative zearalenol glucuronide (modified cosine similarity, 0.345). **e** Comparison of ochratoxin A (OTA) and putative ochratoxin α (OTα), retained through shared OTA-derived anchors despite a modified cosine similarity of 0.646.

The anchoring step was successfully performed for the ZEN seed, generating 27 final seed anchors (Supplementary Fig. 3). Projection of these anchors onto the local feature list retained 95 of 2,343 features (4.1%). Subsequent interpretation yielded phase I/II transformation matches for 28 of these features (29.5%; Fig. 3b). Among these 28 features, 19 (67.9%) showed a modified cosine similarity below 0.70 to ZEN, indicating that many would not form a direct edge to the seed by a conventional spectral-similarity threshold alone (Supplementary Fig. 4). Representative candidates, together with their transformation matches, mass differences, and modified cosine similarities are summarized in Supplementary Table 1.

Among the 28 features with phase I/II transformation matches, one showed a neutral mass difference of 2.01565 Da from ZEN, consistent with hydrogenation (+H□), and was interpreted as a putative zearalenol. Its modified cosine similarity to ZEN was 0.350 despite this small mass difference (Fig. 3c, upper panel). Thus, the putative reduction product retained sufficient ZEN-derived anchors to meet the projection criterion notwithstanding its low whole-spectrum similarity to the parent.

A similar pattern was observed for glucuronide conjugates. Putative zearalenone glucuronide showed relatively high modified cosine similarity of 0.872 to ZEN. In contrast, a feature consistent with hydrogenation followed by glucuronidation, interpreted as putative zearalenol glucuronide, showed a modified cosine similarity of 0.090 to ZEN (Fig. 3c, lower panel). The modified cosine similarity between the two putative glucuronides was 0.345 (Fig. 3d). At the modified cosine threshold of 0.70, putative zearalenone glucuronide appeared in the ZEN-containing molecular-network component, whereas putative zearalenol glucuronide appeared in a separate component (Supplementary Fig. 4). Both features met the ANCHor projection criterion. Their mass differences from ZEN matched glucuronidation and hydrogenation followed by glucuronidation, respectively.

We further applied ANCHor to ochratoxin A (OTA)-treated data within the same spheroid-based mycotoxin metabolism framework. Using OTA as the seed, ANCHor retained a feature that was 147.0684 Da lower in neutral mass than OTA. This mass shift was not covered by common phase I/II transformation rules, but was consistent with the removal of the phenylalanine moiety from OTA to form putative ochratoxin α (OTα). The modified cosine similarity between OTA and the putative OTα feature was 0.646, below the 0.70 threshold (Fig. 3e). Nonetheless, the feature met the ANCHor projection criterion through shared OTA-derived anchors, illustrating the limitation of relying on phase I/II rules alone for interpretation. For such relationships, ANCHor also evaluates KEGG reaction-derived mass shifts as a secondary interpretation layer. KEGG-derived matches were treated as reaction-based hypotheses only when supported by both shared-anchor evidence and mass-difference agreement, rather than as standalone annotation evidence.

### Interpretation of ALD-associated fecal unknown features

We next evaluated whether ANCHor could support the annotation of library-unmatched features by using an unknown feature as the seed, such that annotated features retained by projection served as chemical reference points for the seed. Fecal samples from healthy controls (HC; n = 5) and individuals with alcohol-associated liver disease (ALD; n = 5) were analyzed by untargeted LC–MS/MS (Fig. 4a). To minimize the influence of analytical variation, only features that passed blank filtering and had a pooled-QC RSD below 5% were included in subsequent analyses. Among these, features that lacked a spectral-library match and differed significantly between the two groups were carried forward as potential seeds.

**Fig. 4.**
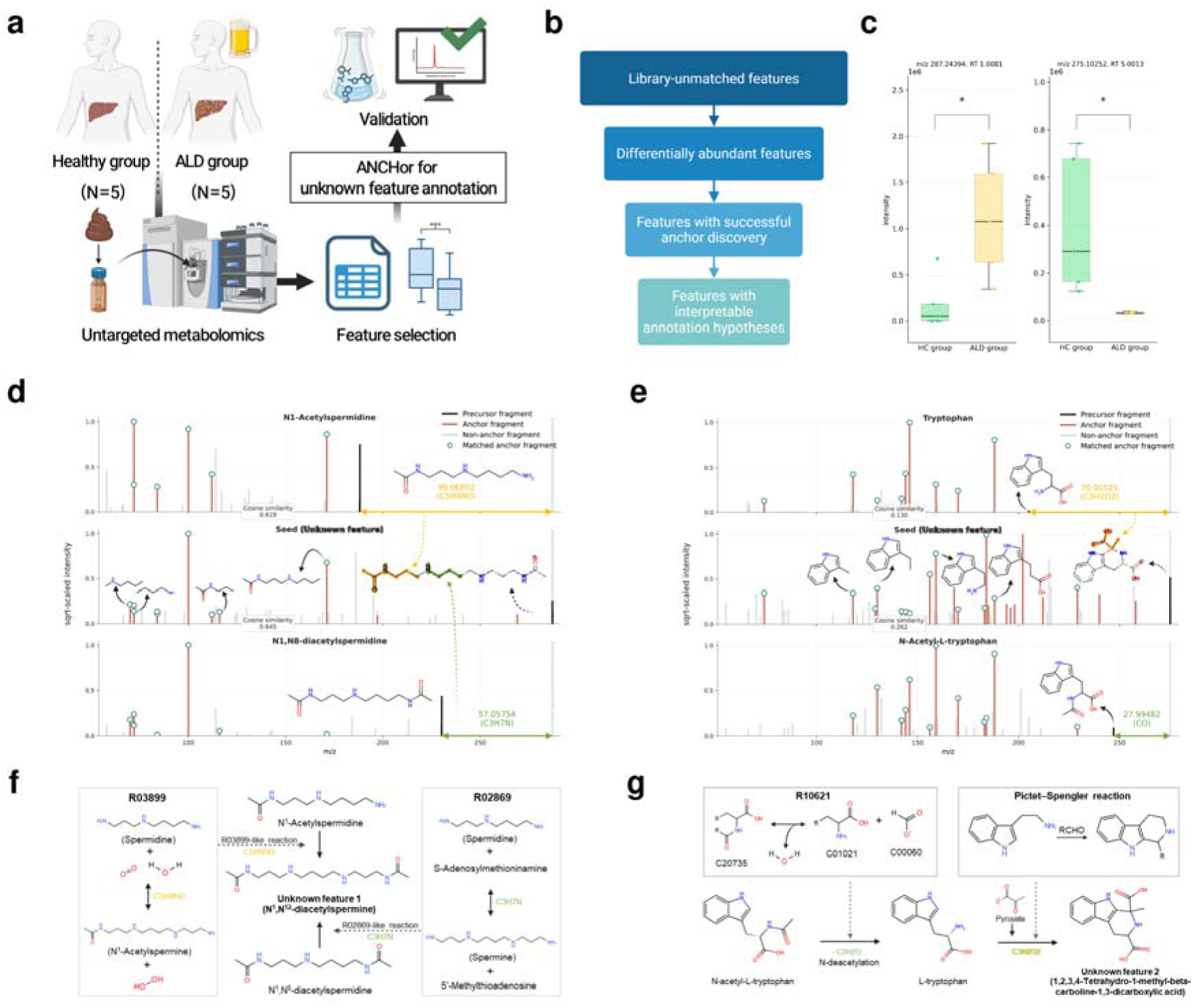
ANCHor-assisted interpretation of ALD-associated fecal features. **a** Workflow for fecal LC–MS/MS analysis and ANCHor-based interpretation of library-unmatched features in healthy controls (HC) and individuals with alcohol-associated liver disease (ALD). **b** Prioritization scheme for selecting library-unmatched features for detailed interpretation. **c** Abundance differences for the two selected unknown features between HC and ALD groups (n = 5 individuals per group; two-sided Mann–Whitney U test; *P < 0.05). **d, e** ANCHor-supported MS/MS comparisons with related annotated features. Red peaks indicate seed anchors, circles indicate matched anchors, and arrows denote precursor mass differences. **f, g** Reaction information supporting the proposed polyaminerelated and tryptophan–pyruvate-derived annotations, respectively. In c, boxes span the 25th–75th percentiles, centre lines indicate medians, and whiskers extend to the most extreme observations within 1.5 times the interquartile range. Dots represent individual participants.

Among the significant library-unmatched features, 245 had acquired MS/MS spectra and were evaluated as seeds. Final seed anchors were generated for 192 of these features. For 148 seeds, projection retained at least one annotated feature, and subsequent interpretation yielded plausible chemical relationships based on neutral mass differences (Fig. 4b). Two library-unmatched features were examined in detail (Fig. 4c).

The first feature was detected at m/z 287.2439 with a retention time of 1.008 min and showed higher abundance in the ALD group than in the HC group (Fig. 4c, left). This feature formed a small molecular subnetwork with three neighboring features, none of which carried a spectral-library annotation (Supplementary Fig. 5, left). By contrast, ANCHor linked this unknown seed to two annotated polyamine-related features, N¹-acetylspermidine and N¹,N□-diacetylspermidine. These features shared 7 and 6 of the 15 seed anchors (47% and 40%), despite modified cosine similarities to the seed of 0.619 and 0.645, below the 0.70 threshold used here for molecular-network edge (Fig. 4d).

Relative to N¹-acetylspermidine, the seed was 99.0680 Da higher in neutral mass, corresponding to ΔC□H□NO. This difference matches the net change from N¹,N¹²-diacetylspermine to N¹-acetylspermidine in the N¹-acetylpolyamine oxidase reaction, which also produces 3-acetamidopropanal (KEGG EC 1.5.3.13; Fig. 4f). Relative to N¹,N□-diacetylspermidine, the seed was 57.0575 Da higher in neutral mass, corresponding to ΔC□H□N. This difference matches the addition of 3-aminopropyl group in the spermidine-to-spermine conversion represented by KEGG reaction R02869. Although R02869 does not directly describe the corresponding acetylated pair, it provided a reaction-class precedent for interpreting the observed mass difference. The mass differences from the two reference features supported assignment of the seed to the acetylated polyamine class, and the N¹-acetylpolyamine oxidase reaction further narrowed the hypothesis to N¹,N¹²-diacetylspermine.

To test this hypothesis, an authentic N¹,N¹²-diacetylspermine standard was analyzed by LC– MS/MS. The standard matched the fecal feature in retention time, but the initial MS/MS spectra differed, with a high-abundance fragment at m/z 186.076 observed only in the fecal spectrum. This additional fragment was attributed to co-isolation interference from a co-eluting ion within the MS/MS isolation window (0.4 Da). Upon reanalysis with a different chromatographic column, the fecal feature and authentic standard showed matching retention times and MS/MS fragmentation patterns (Supplementary Fig. 6, lower panel). The fecal unknown feature was thus confirmed as N¹,N¹²-diacetylspermine, the compound proposed by ANCHor from shared-anchor and mass-difference evidence alone.

Importantly, the ANCHor annotation hypothesis was generated from initial interference-affected fecal spectrum, before the co-isolating ion was recognized. Nevertheless, the interference-derived peak at m/z 186.076 was not retained as a final seed anchor (Supplementary Fig. 7). The anchor-selection criteria require recurrence among seed-neighbor spectra and rarity in background spectra rather than intensity in the seed spectrum, and the co-isolated fragment therefore did not enter the shared-anchor evidence.

The second feature was detected at m/z 275.1025 with a retention time of 5.001 min and showed lower abundance in the ALD group (Fig. 4c, right). This feature also formed a small molecular subnetwork, consisting of two neighboring features, neither of which carried a spectral-library annotation (Supplementary Fig. 5, right). By contrast, ANCHor retained two annotated features for this seed, L-tryptophan and N-acetyl-L-tryptophan, which shared 10 and 12 of the 25 seed anchors (40% and 48%) despite modified cosine similarities to the seed of 0.130 and 0.262 (Fig. 4e).

Relative to L-tryptophan, the seed was 70.0053 Da higher in neutral mass, corresponding to ΔC□H□O. This difference matched pyruvate incorporation with loss of H□O. L-tryptophan and pyruvic acid have been reported to form tetrahydro-β-carboline-1,3-dicarboxylic acid diastereomers^29,30^, and the seed was therefore interpreted as a putative tryptophan–pyruvate-derived Pictet–Spengler product, specifically a diastereomer of 1,2,3,4-tetrahydro-1-methyl-β-carboline-1,3-dicarboxylic acid (Fig. 4g). Relative to N-acetyl-L-tryptophan, the seed was 27.9948 Da higher in neutral mass. This difference was consistent with R10621-like N-deacetylation to L-tryptophan, followed by pyruvate incorporation and H□O loss. The mass difference from L-tryptophan therefore formed the basis for the Pictet– Spengler product hypothesis, while the relationship to N-acetyl-L-tryptophan supported a tryptophan-derived origin for the seed.

To test this hypothesis, L-tryptophan and pyruvic acid were incubated at 50 °C, and the resulting reaction mixture was analyzed using the same LC–MS/MS method. A reaction-generated product matched the fecal unknown feature in precursor m/z, retention time, and MS/MS fragmentation pattern (Supplementary Fig. 8). These concordant measurements supported the annotation of the unknown feature as a diastereomer of 1,2,3,4-tetrahydro-1-methyl-β-carboline-1,3-dicarboxylic acid, the structure proposed by ANCHor from shared-anchor and mass-difference evidence.

KEGG reaction information was not used as standalone evidence for compound identity. Rather, it was used to assess whether the observed seed–feature mass differences were consistent with known biochemical relationships. Overall, these cases show that ANCHor can relate selected library-unmatched features to annotated features, and combine shared-anchor and mass-difference evidence into experimentally testable annotation hypotheses.

### Optional model-assisted structural prioritization

In the preceding case studies, structural hypotheses for unknown seeds were manually formulated from annotated features sharing seed anchors and plausible transformations consistent with the observed mass differences. These references can support different structural hypotheses for the same seed. We therefore implemented an optional Transformer-based candidate-prioritization module that integrates evidence from annotated features retained after ANCHor projection (Supplementary Fig. 9). Training combined curated reaction-supported cases, including variants with controlled distractors, with rule-derived pseudo-labelled cases (Supplementary Fig. 9a). During application, the trained model used structural similarity and local transformation evidence to calculate candidate–reference compatibility scores and relative reference weights, which were combined to rank candidate two-dimensional structures sharing the reconstructed seed formula (Supplementary Fig. 9b).

In the held-out test set of 356 targets, the correct two-dimensional structure ranked first for 80.34% of targets (286/356) and within the top five for 98.03% (349/356), with a mean reciprocal rank of 0.87723. Candidate scores were averaged across the available reference-set variants before ranking, and each target contributed once to these metrics.

With the selected model parameters fixed, we retrospectively applied the module to the two fecal cases described above. For the tryptophan–pyruvate-derived feature, the β-carboline two-dimensional proxy supported by reaction-product analysis ranked first among 85 naturally retrieved candidates. For the N¹,N¹²-diacetylspermine case, the authentic-standard-supported two-dimensional structure ranked first among seven candidates.

## Discussion

Our results show that structurally or biochemically related molecules can retain shared MS/MS fragments even when overall spectral similarity is low. ANCHor leverages this fragment-level information to evaluate possible seed-feature relationships that may not be apparent from overall MS/MS similarity alone. Seed-neighbor spectra retrieved from the DreaMS atlas were used to assess the recurrence of fragments observed in the seed and their rarity in background spectra. Fragments meeting these criteria were defined as anchors. These anchors were projected onto the dataset to identify features sharing seed anchors. Mass differences and transformation knowledge were then used to interpret their possible relationships to the seed and generate testable annotation hypotheses.

Unknown metabolite interpretation has developed along several directions, including spectral networking (FBMN)^15^, spectral-library analogue search (MS2Query)^31^, recurring fragment-discovery (MS2LDA/Mass2Motif)^32^, and user-defined pattern querying (MassQL)^33^. Other approaches use reaction knowledge to guide annotation (MetDNA3^34^ and BAM^35^) or generate candidate structures (MSNovelist^36^ and MetGenX^37^).

ANCHor complements these approaches with a seed-centered workflow that combines atlas-supported fragment anchors with mass-difference evidence to identify local features related to an unknown seed. The same workflow can prioritize putative transformation products of a known parent compound or retrieve annotated features as chemical references for an unknown seed, without requiring prior annotation of the seed.

The methionine panel and mycotoxin cases provided known-seed examples of this workflow, showing that ANCHor retained chemically plausible features that were not captured by MS/MS similarity-based networking. In particular, some putative reduction and conjugation products appeared in network components separate from the seed. FBMN may recover some of these relationships at a lower similarity threshold, but it may also include less-related spectra. ANCHor instead applies a separate anchor-sharing criterion to the selected seed.

The most intense peaks are not necessarily the most informative for interpreting chemical relationships. In the methionine pseudo-spectrum, ANCHor excluded a relatively intense peak that was frequent in BG1 despite recurring among seed-neighbor spectra, whereas it retained lower-intensity fragments meeting the recurrence and background-specificity criteria. In the polyamine case, ANCHor did not select the intense signal at m/z 186.076 in the initial fecal MS/MS spectrum as an anchor. ANCHor generated an annotation hypothesis for N¹,N¹²-diacetylspermine using shared-anchor and mass-difference evidence. Subsequent analysis using a different chromatographic separation indicated that this signal likely originated from co-isolation interference.

Both fecal unknowns in Fig. 4 belonged to small FBMN subnetworks, but their neighboring features lacked library annotations, limiting the available structural context. ANCHor linked these unknowns to annotated polyamine-or tryptophan-related features outside their original subnetworks. Mass differences between each unknown and multiple annotated references suggested plausible chemical transformations. In both cases, ANCHor-derived relationships led to structural hypotheses that were subsequently evaluated using an authentic standard or a reaction-generated product.

Multiple annotated seed-related features can inform the structure of an unknown seed, but their contributions may differ. In the preceding case studies, we integrated the information from these references manually to formulate structural hypotheses. To assist this step, the optional candidate-prioritization model uses learned relative weights to combine reference information and rank 2D candidate structures. Structure-based testing and retrospective application to the two fecal cases suggest that the model can help prioritize candidates for further evaluation. The resulting weights and ranks should not be interpreted as probabilities that individual relationships or compound identities are correct.

The candidate-prioritization model requires successful anchor generation. In the authentic-standard benchmark (Fig. 2a), lipids and lipid-like molecules had the highest anchor-generation rate among the five most represented superclasses (81.7%). Rates varied across lipid-related categories. At the class level, steroids and steroid derivatives reached 94.3%. Among lipid-related subclasses, anchor-generation rates were 100% for fatty acid esters and 65.7% for fatty acids and conjugates. Non-lipid purine nucleotides also showed a high rate of 90.9%. Uneven representation across chemical categories limits direct comparisons. Fatty acid esters had a median of seven seed peaks, lower than the median for fatty acids and conjugates. ANCHor evaluates seed peaks individually, so more peaks may offer more opportunities for anchor selection, but peak count alone is not sufficient for successful anchoring. ANCHor selection requires at least one seed peak that recurs among seed-neighbor spectra, remains rare in background spectra, and satisfies the statistical selection criteria.

The current implementation depends on the DreaMS model and atlas coverage, and is restricted to positive ionization mode. ANCHor may fail to generate anchors when too few neighbors are retrieved or no seed peak meets the selection criteria. Such failure does not imply the absence of chemical relationships. Even when ANCHor generates anchors, the requirement for at least two shared anchors can limit projection, particularly for seeds with few anchors. These projection criteria were predefined and applied consistently throughout this study. For other applications, users may adjust these criteria, but should assess the effects on the recovery of related features and nonspecific matching. Incorrect adduct assumptions can also affect mass-difference interpretation.

In summary, ANCHor supports the structural interpretation of library-unmatched features by relating them to known metabolites and plausible chemical transformations. This approach provides chemical context for unknowns that remain difficult to interpret through conventional spectral comparisons and guides the development and experimental evaluation of structural hypotheses.

## Methods

### HepG2 spheroid culture and metabolite extraction

HepG2 cells (American Type Culture Collection) were maintained in Minimum Essential Medium Eagle supplemented with 10% fetal bovine serum (Biowest), 1% L-glutamine (Gibco) and 1% penicillin–streptomycin (Gibco). Spheroids were generated in 96-well ultra-low-attachment U-bottom plates at 5 × 10³ cells per well following the previously described culture conditions^38^. After seeding, plates were centrifuged at 1,250 rpm for 10 min and incubated at 37 °C under 5% CO, with medium replacement every 48 h. Zearalenone (ZEN; TargetMol) and ochratoxin A (OTA; Sigma-Aldrich) stock solutions were prepared in dimethyl sulfoxide and stored at −20 °C. Following an initial 3-day spheroid formation period, spheroids were exposed separately to 50 μM ZEN or 10 μM OTA for 7 days, corresponding to the high-concentration conditions reported for the HepG2 3D model in Supplementary Table S2 of Moon et al.^38^. Untreated spheroids served as controls. Three biological replicates per condition were included in the ANCHor analysis.

For extraction of extracellular metabolites, 400 μL of conditioned medium was combined with 1.2 mL of isopropanol:methanol (1:1, v/v). The mixture was vortexed for 10 s, sonicated for 10 min and centrifuged at 13,200 rpm for 10 min at 4 °C. A 1.4-mL aliquot of the supernatant was transferred to a fresh tube and dried in a speed vacuum concentrator. The dried extract was reconstituted in 50 μL of water:methanol (1:1, v/v) before LC–MS analysis^38^.

### Human participants and ethics

All patients met histological or established clinical criteria for cirrhosis and provided written informed consent. The study protocol complied with the ethical principles of the 1975 Declaration of Helsinki and was approved before study initiation by the Institutional Review Board of Hallym University Chuncheon Sacred Heart Hospital (approval nos. 2016_11_134 and 2019_09_001).

### Fecal metabolite extraction

Fecal samples (100 mg) were transferred to 2.0-mL round-bottom Eppendorf tubes and thawed at 4 °C. A stainless-steel bead and 1.4 mL of cold methanol:isopropanol:water (3:3:2, v/v/v) were added to each sample. After vortexing for 10 s, samples were homogenized in a mixer mill at 25 Hz for 90 s, sonicated on ice for 10 min and incubated on ice for a further 30 min39. Extracts were centrifuged at 13,200 rpm for 10 min at 4 °C, and 600 μL of the supernatant was transferred to a fresh 1.5-mL tube. This aliquot was dried completely in a speed vacuum concentrator and stored at −80 °C until analysis. Before LC–MS analysis, dried extracts were reconstituted in 100 μL of acetonitrile:water:methanol (70:25:5, v/v/v), sonicated on ice for 10 min and transferred to polytetrafluoroethylene tubes. Following centrifugation at 9,000 rpm for 5 min at 4 °C, the supernatants were transferred to LC–MS vials^40^.

### Reversed-phase liquid chromatography

Untargeted metabolomic profiling was performed using reversed-phase LC–MS. Separations used a Waters ACQUITY UPLC BEH C18 column (100 × 2.1 mm, 1.7-μm particles) fitted with a BEH C18 VanGuard pre-column (5.0 × 2.1 mm, 1.7-μm particles). The flow rate was 0.30 mL min□¹, the injection volume was 2 μL and the column temperature was 40 °C.

On the Thermo Scientific UltiMate 3000 UPLC system, mobile phase A consisted of water containing 0.1% formic acid, and mobile phase B was acetonitrile. The gradient was maintained at 0.5% B for 0–0.1 min, increased linearly to 80% B at 10 min and then to 99.5% B at 10.1 min. This composition was held until 12 min, followed by re-equilibration at 0.5% B from 12.1 to 15 min.

Spheroid samples were analysed on a Thermo Scientific Vanquish UHPLC system using the same column dimensions, flow rate, injection volume and column temperature. Mobile phases A and B were water and acetonitrile, respectively, both containing 0.1% formic acid. The 15-min gradient began at 0.5% B for 0.1 min, increased to 80% B at 10 min, and was followed by a 2-min hold at 99.5% B and a 3-min re-equilibration at 0.5% B^38^.

### HILIC analysis for confirmatory comparison

HILIC was used for confirmatory comparison of the fecal feature with the authentic N¹,N¹²-diacetylspermine standard. Sample extraction and handling followed the procedures described above. Dried extracts were reconstituted in acetonitrile:distilled water:methanol (70:25:5, v/v/v). Separations were performed on a Waters ACQUITY UPLC BEH Amide column (150 × 2.1 mm, 1.7-μm particles) with a matching guard column at 45 °C and a flow rate of 0.40 mL min ¹. Mobile phase A was water containing 10 mM ammonium formate and 0.125% formic acid. Mobile phase B was acetonitrile:water (95:5, v/v) containing the same additives. The gradient was 0–2 min, 100% B; 2–7.7 min, 100–70% B; 7.7–9.5 min, 70–40% B; 9.5–10.25 min, 40–30% B; and 10.25–17 min, 100% B.

### Mass spectrometry

The UltiMate 3000 system was coupled to a Q Exactive Plus Orbitrap mass spectrometer (Thermo Fisher Scientific) equipped with a heated electrospray ionization source. Samples were injected in randomized order. Full-scan spectra were acquired over m/z 80–1,200 at a resolution of 70,000 FWHM at m/z 200, with an automatic gain control (AGC) target of 1 × 10□. Data-dependent MS/MS spectra were collected at a resolution of 17,500 using stepped normalized collision energies of 30, 40 and 50. The system was operated using Q Exactive Tune and Xcalibur 4.0.

For spheroid samples, the Vanquish UHPLC was coupled to a Q Exactive Focus Orbitrap mass spectrometer (Thermo Fisher Scientific). Data were acquired in positive and negative ionization modes over m/z 80–1,200. Spray voltages were 3.8 kV and 3.5 kV in positive and negative modes, respectively. The capillary and probe heater temperatures were 320 °C and 300 °C, and the sheath and auxiliary gas settings were 40 and 10. Full-scan and data-dependent MS/MS resolutions were 70,000 and 17,500, respectively. The three most intense eligible precursor ions were selected for MS/MS with an isolation window of 0.4 m/z and normalized collision energies of 30, 40 and 50. Acquisition used an AGC target of 1 × 10□, a maximum injection time of 50 ms, an intensity threshold of 2 × 10³, an apex trigger of 3–6 s and a dynamic exclusion of 5 s. The instrument was calibrated weekly using positive- and negative-ion calibration solutions and controlled with Q Exactive Tune and Xcalibur 4.0^38^. Positive-ion MS/MS spectra were used for ANCHor analysis.

### LC–MS data processing

Raw LC–MS files were processed in MZmine 4^41^. Mass detection used a noise factor of 5. Chromatogram construction required at least five consecutive scans exceeding an intensity threshold of 1 × 10□, with an m/z tolerance of 5 ppm. Feature lists were aligned using retention-time and m/z tolerances of 0.5 min and 5 ppm, respectively. Spectral library matching used precursor and fragment mass tolerances of 5 ppm and 10 ppm. Features with intensities below five times the corresponding blank signal were removed to reduce background interference. Feature tables and associated MS/MS spectra were exported for ANCHor analysis.

Pooled quality-control (QC) samples were prepared by pooling aliquots from all fecal samples and were analysed five times. Features with a pooled-QC relative standard deviation below 5% were retained for subsequent fecal analyses.

Feature-based molecular networking (FBMN)^15^ was performed using default parameters.

### Statistical analysis

Differences in feature intensities between the healthy control and alcohol-associated liver disease groups were assessed using two-sided Mann–Whitney U tests. *P*-values were adjusted using the Benjamini–Hochberg procedure, and an adjusted *P*-value < 0.05 was considered statistically significant. For the two selected features shown in Fig. 4c, unadjusted two-sided *P*-values are displayed, with \**P* < 0.05 indicating statistical significance.

### Input data

LC–MS/MS data were preprocessed using feature detection software such as MZmine 4. ANCHor requires two input files: (i) a feature table containing feature IDs, precursor m/z values, retention times, and sample intensities, and (ii) an MGF file containing MS/MS spectra for the corresponding features. A feature of interest is specified by its feature ID and used as the seed for downstream analysis.

### DreaMS-based atlas retrieval

DreaMS^42^ was used to represent MS/MS spectra as embeddings generated by a model trained on large-scale unannotated spectra. This representation enables comparison of fragmentation patterns related to molecular structure beyond direct peak overlap. The seed MS/MS spectrum was encoded into a DreaMS embedding using up to the 100 most intense fragment peaks. Atlas spectra were first prefiltered using a 10 ppm precursor m/z tolerance relative to the seed precursor m/z.

The prefiltered atlas spectra were then ranked by DreaMS similarity, defined as the cosine similarity between each precomputed atlas embedding and the seed embedding. Spectra with DreaMS similarity ≥ 0.70 were retrieved as seed neighbors. These atlas-derived seed neighbors formed the reference population for anchor discovery. The experimental seed was excluded from occurrence counts and enrichment calculations. Spectra with DreaMS similarity ≤ 0.30 were assigned to background set 1, and 1,000 spectra randomly sampled from the atlas were used as background set 2. Background set 1 therefore provided a precursor-m/z-matched contrast for anchor selection, whereas background set 2 provided an atlas-wide measure of fragment rarity. The 0.30 was set as the complement of the 0.70 similarity criterion used to define seed neighbors, and was intended to exclude spectra of intermediate similarity from the contrast rather than to serve as an anchor-selection criterion.

### Seed anchor selection (anchoring)

Seed anchors were selected from fragment centroids using DreaMS-retrieved seed-neighbor spectra and two background sets. Anchor fragments were defined as seed-associated fragment signals that were present in the seed spectrum, recurrent among seed-neighbor spectra, and infrequent in background spectra.

From the preprocessed MS/MS peak lists, fragment peaks in the seed-neighbor spectra were aligned into m/z centroids, and the occurrence frequency of each centroid was calculated across the seed-neighbor and background spectra. For each centroid, enrichment among seed-neighbor spectra relative to background set 1 was assessed using a one-sided hypergeometric test, followed by Benjamini–Hochberg false-discovery-rate (FDR) correction.

Final anchor fragments were defined as centroids that were present in the seed spectrum, sufficiently recurrent among seed-neighbor spectra, and infrequent in both background sets. Specifically, each centroid was required to occur in at least 30% of the seed-neighbor spectra while occurring in less than 15% of background set 1 and less than 20% of background set 2. Centroids below m/z 80 additionally required an FDR-adjusted q-value ≤ 0.05 and a natural-log odds ratio ≥ 0.50 for enrichment relative to background set 1. These two enrichment thresholds were not required for centroids at m/z ≥ 80. The core-support and background-frequency criteria had to be satisfied in at least 80% of bootstrap resampling iterations; the additional low-mass enrichment criteria were evaluated using the original populations.

### Anchor projection onto the local feature list (projection)

The selected seed anchors were projected onto all MS/MS spectra in the local feature list to identify features containing seed anchors. For each feature, the number of fragments matching the seed anchors was calculated using a mass tolerance of 10 ppm or 0.010 Da.

Features sharing a sufficient number and proportion of seed anchors were retained as projection-retained features. Specifically, the minimum required number of shared anchors was set to max(2, ceil(0.20 × N_seed)), where N_seed denotes the number of seed anchors used for projection. A feature was retained if the number of shared anchors met this threshold and if at least 20% of its processed spectral peaks matched seed anchors. Projection-retained features were then forwarded to transformation-based candidate annotation.

### Transformation-based candidate annotation (interpretation)

Projection-retained features were evaluated using knowledge-based transformation rules to infer possible chemical relationships to the seed. For each seed–feature pair, neutral masses were calculated from the assumed ion forms, and the observed neutral mass difference was compared with mass shifts corresponding to phase I, phase II, and KEGG reaction-derived transformations^43^.

Features with precursor m/z values nearly identical to that of the seed were classified as isomer candidates. The remaining features were matched against the transformation rule table. When both phase I/II and KEGG reaction-derived transformations were possible, phase I/II transformations were prioritized. A transformation match was accepted when the observed neutral mass difference fell within the tolerance window for the expected mass shift.

### Model-assisted structural prioritization

Known structures were treated as hidden targets and paired with positive references from curated reaction-supported relationships. Controlled distractors represented incorrect proposed links to otherwise correctly identified reference structures. Additional training cases were generated using local transformation rules extracted exclusively from curated training relationships and were treated as down-weighted pseudo-labels. The combined training set contained 10,345 targets and 11,689 reference-set variants; validation and test sets contained 227 and 356 curated targets, respectively. Evaluation target components were excluded from all training roles, although candidate alternatives were not globally disjoint across splits.

Candidate two-dimensional structures matching the reconstructed seed formula were retrieved from a frozen PubChem-derived registry^44^. For each candidate–reference pair, the model received 12 numerical features summarizing structural similarity, local transformation evidence and candidate context. Structural similarities were calculated using radius-2 Morgan count fingerprints of size 2,048^45^. A shared encoder and a two-layer, four-head Transformer^46^ without positional encoding learned candidate–reference compatibility scores and relative reference weights, which were combined to rank candidates. Training combined candidate-ranking and auxiliary reference-role losses, and checkpoint selection used validation target-level mean reciprocal rank. Detailed procedures are provided in Supplementary Methods.

### Reference standard and reaction-product comparisons

An authentic N¹,N¹²-diacetylspermine standard (catalogue no. 91423-100MG) was used for LC–MS/MS comparison with the fecal feature.

For preparation of the tryptophan–pyruvic acid reaction mixture, 2.04 mg of L-tryptophan was combined with 100 μL of a 1 M pyruvic acid stock solution, 10 μL of a 10% formic acid stock solution and 890 μL of LC–MS-grade water, giving a total volume of 1 mL. The mixture was incubated at 50 °C for approximately 10 h and subsequently analysed by LC– MS/MS.

## Data availability

The raw LC–MS/MS data supporting the findings of this study have been deposited in Zenodo and are available at https://doi.org/10.5281/zenodo.23553283.

## Code availability

The source code for ANCHor is hosted at GitHub (https://github.com/SNUFML/ANCHor). The version used for the analyses reported in this study (v0.5.0rc2) will be archived in Zenodo under DOI 10.5281/zenodo.23555440. The repository will be made publicly accessible upon publication.

## Acknowledgements

This work was supported by the Advanced GPU Utilization Support Program and National Research Foundation of Korea (NRF) grants (Nos. RS-2023-00273634 and RS-2022-NR068419), funded by the Ministry of Science and ICT (MSIT), Government of the Republic of Korea.

## Author contributions

Y.J.P. and D.Y.L. conceived the study and developed the methodology. Y.J.P. performed the metabolomics experiments and computational analyses, developed the software, and prepared the figures. J.H.M. performed the cell-based experiments using the HepG2 spheroid model. K.T.S. contributed clinical samples and clinical information. D.Y.L. supervised the study. Y.J.P. wrote the original manuscript draft. All authors reviewed and edited the manuscript and approved the final version.

