## Supplementary Information for "Fragment anchoring for metabolite annotation beyond whole-spectrum similarity"

### Supplementary Methods

#### ANCHor spectral relationship analysis

#### Inputs and spectral preprocessing

ANCHor used a local feature list containing feature identifiers, precursor m/z, retention times, sample intensities and available annotations, together with feature-linked MS/MS spectra in MGF format, a spectral atlas and knowledge-based transformation rules. The implementation supported singly charged positive ions and required an experimental MS/MS spectrum for each seed.

Peaks with nonfinite m/z or intensity values or nonpositive intensities were removed. Sorted peaks were grouped sequentially by comparison with the current group median, retaining the most intense peak per group. Mass tolerances were the greater of 10 ppm and 0.005 Da for anchoring, or 0.010 Da for projection. A higher-mass peak was removed when a peak 1.003355 Da below it, within 0.01 Da, had equal or greater intensity. Seed embeddings were generated using pretrained DreaMS^1^, without fine-tuning, from up to the 100 most intense peaks.

#### Ion forms and atlas retrieval

Known mode assumed [M+H]⁺ for both seed and experimental features; this designation did not imply confirmed structural identity. Unknown mode considered [M+H]⁺, [M+Na]⁺, [M+NH₄]⁺, [M+K]⁺ and [M]⁺, with hydrated or dehydrated protonated forms and protonated or sodiated dimers additionally considered when supported by coeluting features. Neutral monomer mass was calculated by subtracting the ion-form mass shift from the observed precursor m/z and dividing by molecular multiplicity.

Support was sought from features within 0.01 min of the seed, excluding one- or two-isotope spacings. Neutral masses inferred under compatible ion forms were compared using the combined tolerance below, where τ is the greater of 10 ppm and 0.0005 Da, k is molecular multiplicity, and s and c denote the seed and coeluting feature. At most one alternative seed hypothesis was tested after failure or insufficient anchor or projection evidence.

$$\tau_{pair} = \sqrt{\left( \frac{\tau_{s}}{k_{s}} \right)^{2} + \left( \frac{\tau_{c}}{k_{c}} \right)^{2}}$$

Atlas spectra were prefiltered within ±10 ppm of the observed seed precursor m/z. Retrieved atlas spectra with DreaMS embedding cosine similarity ≥0.70 were termed seed-neighbor spectra. The experimental seed was excluded from this reference population and from occurrence counts and enrichment calculations. Retrieved spectra with similarity ≤0.30 formed background set 1 (BG1). Intermediate-similarity spectra entered neither group. Background set 2 (BG2) contained up to 1,000 spectra sampled uniformly without replacement from eligible atlas entries, excluding all mass-retrieved spectra.

#### Seed anchor selection

Seed anchors were fragments present in the seed spectrum, recurrent among seed-neighbor spectra and rare in the background sets. Candidate centroids were median masses of grouped fragments from seed-neighbor spectra. Binary fragment occurrence was recorded across seed-neighbor and background spectra, and centroids were matched to fragments in the experimental seed.

A centroid had to match a seed fragment, occur in at least 30% of seed-neighbor spectra, and occur in <15% of BG1 and <20% of BG2 spectra. Enrichment among seed-neighbor spectra relative to BG1 was evaluated using a one-sided hypergeometric test, with Benjamini–Hochberg adjustment across all candidate centroids. Only centroids below m/z 80 additionally required q ≤ 0.05 and a natural-log odds ratio ≥ 0.50; these two thresholds were not required for centroids at m/z ≥ 80. BG1 and BG2 were independently resampled with replacement ten times, keeping core support fixed. The core-support and background-frequency criteria had to be satisfied in at least 80% of iterations. The additional low-mass enrichment criteria were evaluated using the original populations. Final anchor masses were the corresponding observed seed-fragment masses.

#### Projection and transformation interpretation

Seed anchors were projected onto the local feature list by one-to-one peak assignment, maximizing distinct matches before minimizing summed tolerance-normalized mass error. Let N_seed_ be the number of selected anchors recovered in the processed seed, H the number matched in a feature and N its processed peak count. Projection-retained features satisfied H/N ≥ 0.20 and H ≥ H_min_, where

$$H_{min} = max\left( 2, \left\lceil0.20 N_{seed} \right\rceil\right)$$

The seed was excluded from transformation evaluation. Features matching its observed precursor within the projection tolerance were labelled operational isomer candidates, without establishing identical formulae or structural isomerism. For other features, the observed neutral-mass difference Δ_obs_ (feature minus seed) was compared with each supplied rule mass difference Δ_rule_:

$$|\Delta_{obs} - \Delta_{rule}| \leq max\left( 0.0005, |\Delta_{rule}| \times{10}^{-5} \times\lambda\right) Da$$

Here λ was 2 in known mode and 1 in unknown mode. Unknown mode initially assigned each feature the seed ion form, followed, for eligible monomers, by alternative protonated, sodiated, ammoniated, hydrated-protonated or dehydrated-protonated forms. Within each stage, phase I/II rules preceded KEGG reaction-derived rules. The first successful stage returned up to three matches ranked by support score and absolute mass error. Only individual input rules were evaluated. Support scores ranged from 0 to 2, awarding one point for a retention-time difference ≤0.10 min and one for a precursor-to-fragment or fragment-pair difference consistent with |Δ_rule_| in the higher-precursor spectrum.

#### Unknown mode filtering and spectral comparison

Non-isomer relationships in unknown mode additionally required an anchor-associated precursor loss or shifted fragment pair consistent with the rule mass difference. Both shift directions were tested without peak reuse. Coeluting seed–feature pairs (≤0.01 min) were screened for isotope spacings, common losses (H₂O, NH₃, CO, CO₂ and H₂), alternative adducts and monomer–dimer relationships. Comparisons among transformation-matched features identified potential in-source fragments and alternative adducts. These heuristic filters excluded seed-associated artifacts and retained one representative per ion family using annotation availability and spectral support.

Final unknown-mode reports required projection, transformation and fragment support, artifact exclusion and an existing feature annotation. Matches were grouped by seed, seed ion form, transformation category and name, and mass-difference sign; groups exceeding three hit rows were excluded.

Modified cosine provided supplementary spectral evidence and informed ion-family representative selection. Intensities were square-root transformed, the six most intense peaks per 50 Da window were retained, and peaks within 17 Da of each precursor were removed. Direct and precursor-difference-shifted matches were selected greedily by intensity product without peak reuse and normalized by the intensity-vector norms. Modified cosine was not a general acceptance threshold. Background sampling and bootstrapping used random seed 42. Reported relationships supported annotation hypotheses rather than independently establishing structure or reaction direction.

#### Optional candidate prioritization

An optional module ranked candidate two-dimensional (2D) structures using annotated features retained after ANCHor projection as structural references (Supplementary Fig. 9). Reference identities were assumed correct, but proposed seed–reference links could be incorrect. RDKit^2^ standardized neutral CHNOPS-containing organic parents; connectivity was defined by the first InChIKey block. Seed formulae were reconstructed from reference formulae and signed elemental differences, requiring mass agreement within the greater of 0.005 Da and 20 ppm. Candidates were retrieved by exact formula from a frozen PubChem-derived registry of 899,074 structures^3^, without inserting known answers. Each seed/formula group was ranked separately.

Curated relationships were supported by Rhea^4^ and/or KEGG RCLASS^5^. After filtering, training, validation and test contained 903, 227 and 356 curated targets, respectively. Training additionally included 9,442 rule-derived targets treated as pseudo-labels. Curated targets had up to five positive references and clean or distractor-containing variants; pseudo-labelled targets had up to three references and one clean variant. Distractors were same-formula alternatives to positive references. Incorrect-link labels required complete rule application with either local-pattern mismatch or generation of a competing candidate without the target; unresolved cases were not labelled negative.

The 1,914 local transformation rules, covering 74 elemental differences, were derived only from curated training relationships using maximum-common-substructure correspondence (≥60% of the larger molecule), at most four changed bonds, and radius-1 or radius-2 local environments. Rules had to regenerate the recorded target; products had to be valid, single-component and formula-consistent. Invalid products or more than 64 products indicated incomplete application. Pseudo-label generation required a single valid target without conflicting targets among examined templates. Training excluded rules supported solely by the target’s original component. Components containing evaluation targets were excluded from all training roles, and five reserved structures were excluded from all dataset roles. Validation also excluded test structures; test candidate alternatives could include eligible training or validation structures.

Twelve input features summarized structural similarity, exact transformation-product matches, product specificity, rule availability and application status, product and matching-rule counts, and agreement across references. Similarities used radius-2, 2,048-element Morgan count fingerprints^6^. A two-layer Transformer^7^ implemented in PyTorch^8^ used 64-dimensional representations, four heads and dropout 0.1, with masked padding and no positional encoding. It produced candidate–reference compatibility scores and reference-role outputs. For each reference, the maximum role output across candidates was normalized across references by softmax; candidate scores were the resulting weighted sums of compatibility scores. These weights were relative contributions, not attention matrices or calibrated identification probabilities.

Training combined ranking cross-entropy and reference-role binary cross-entropy (coefficient 0.1). Inverse variant-count weighting equalized target contributions within each source, with curated and pseudo-labelled targets contributing 75% and 25% of the objective. AdamW^9^ used a learning rate of 5 × 10⁻⁴, weight decay of 10⁻⁴, batch size 32 and gradient-norm clipping at 2.0. Training started from fresh initialization with seed 20260818, a maximum of 40 epochs, early-stopping patience of eight and no learning-rate scheduler. Epoch 3 was selected from 11 completed epochs by validation target-level mean reciprocal rank (MRR).

Candidate scores were averaged across selected reference-set variants before ranking, evaluating each target once. Recall@1 and Recall@5 measured the fractions of correct structures ranked first and within five; MRR averaged reciprocal target ranks. Evaluation considered all, clean-only or noisy-only variants; 356 test targets were evaluable overall and 289 had noisy variants. Predictions were saved before comparison with answers. Evaluation remained retrospective because developers knew the application cases and historical evaluation material; it did not establish prospective, scaffold-independent or reaction-family-independent performance. Outputs comprised rankings, relative reference weights, evidence and excluded inputs, with names and PubChem identifiers added after scoring. Predictions were restricted to registry candidates and 2D connectivity.

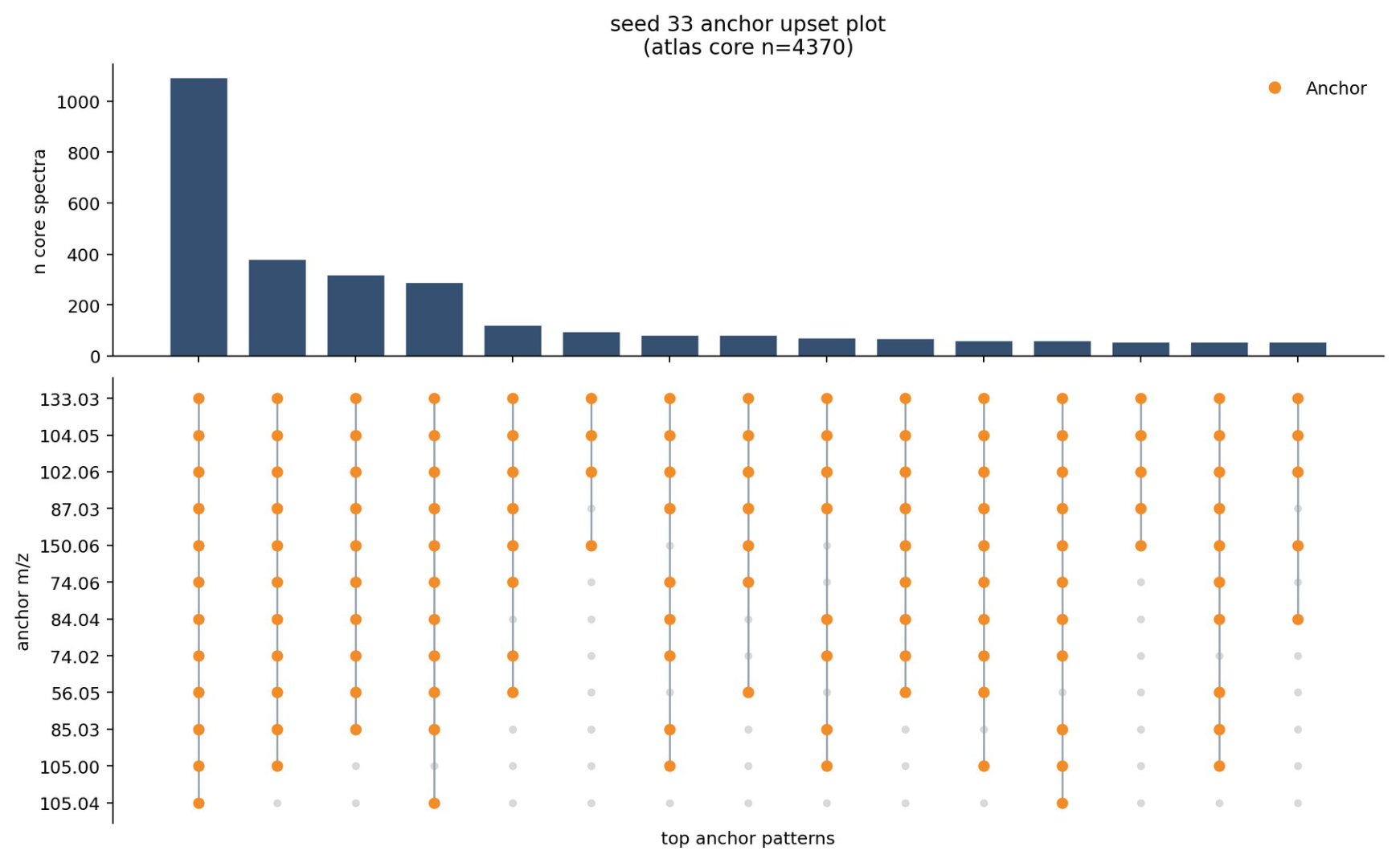

**Supplementary Fig. 1. Occurrence patterns of L-methionine-derived seed anchors.** UpSet plot showing occurrence patterns of the 12 L-methionine seed anchors across atlas reference spectra. Each column represents an anchor combination, and bar heights indicate the number of spectra exhibiting that pattern. Orange dots indicate anchor fragments.

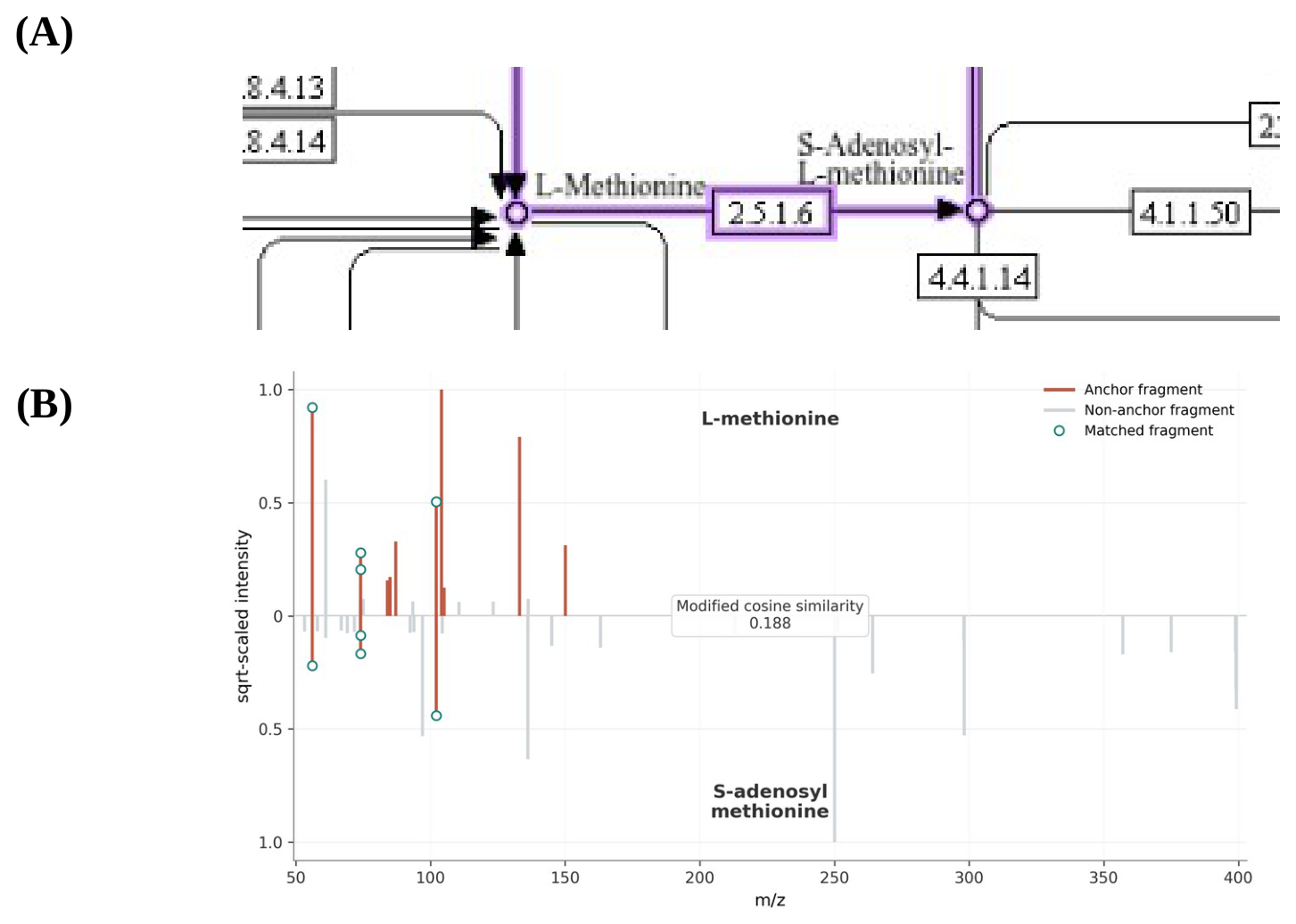

**Supplementary Fig. 2. Spectral similarity and metabolic context of methionine. a** Region of the KEGG cysteine and methionine metabolism pathway showing the direct metabolic connection between L-methionine and S-adenosyl-L-methionine. **b** Mirror plot comparing their MS/MS spectra. The spectra shared four of the 12 L-methionine seed anchors despite a modified cosine similarity of 0.188. Intensities are square-root transformed.

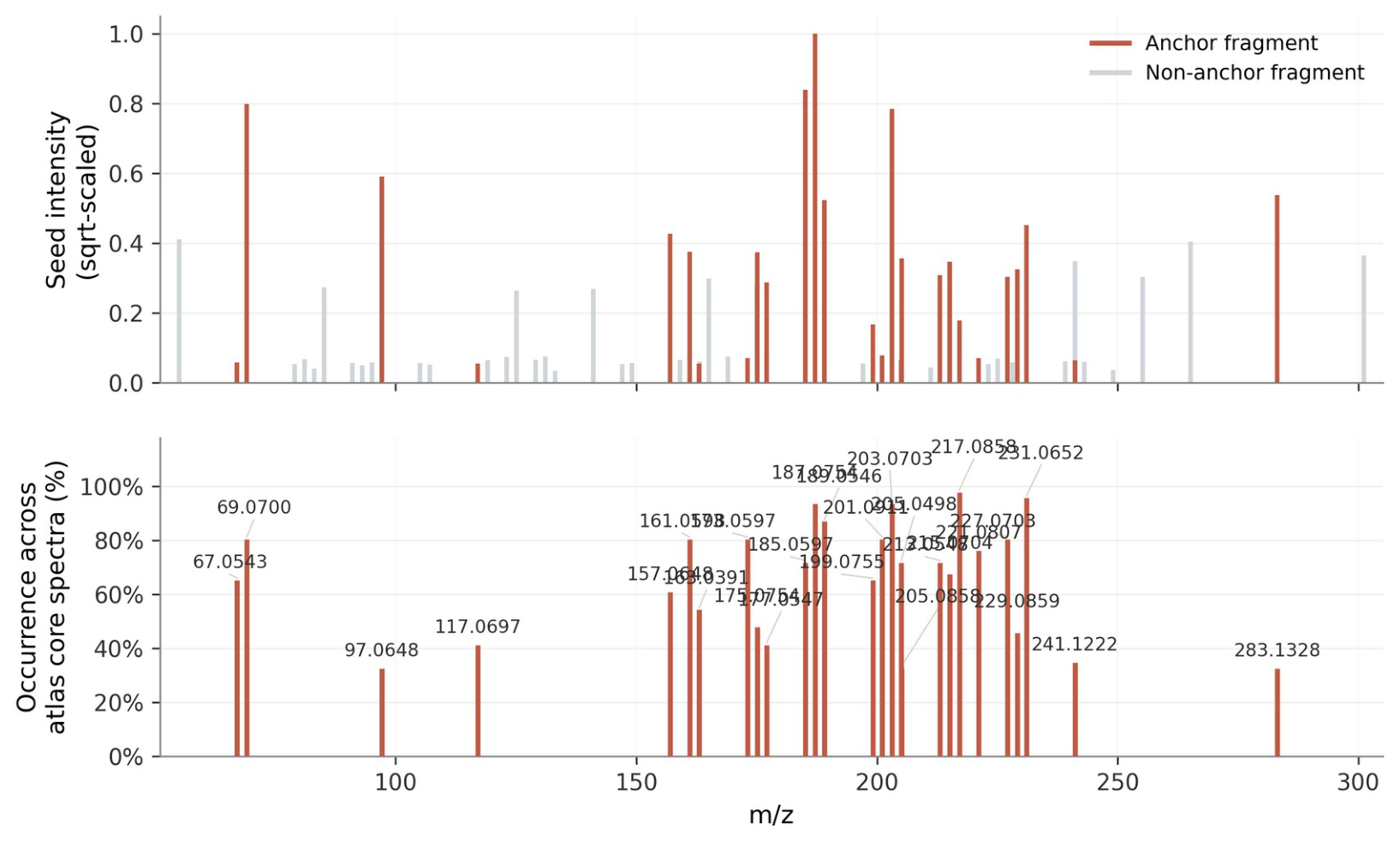

**Supplementary Fig. 3. Atlas-informed selection of zearalenone seed anchors.** Experimental MS/MS spectrum of the zearalenone (ZEN) seed and occurrence frequencies of selected anchors across atlas reference spectra. Experimental intensities are square-root transformed. Highlighted fragments indicate selected anchors, and grey fragments indicate unselected peaks. Atlas-derived peak heights represent fragment occurrence rather than experimental signal intensity.

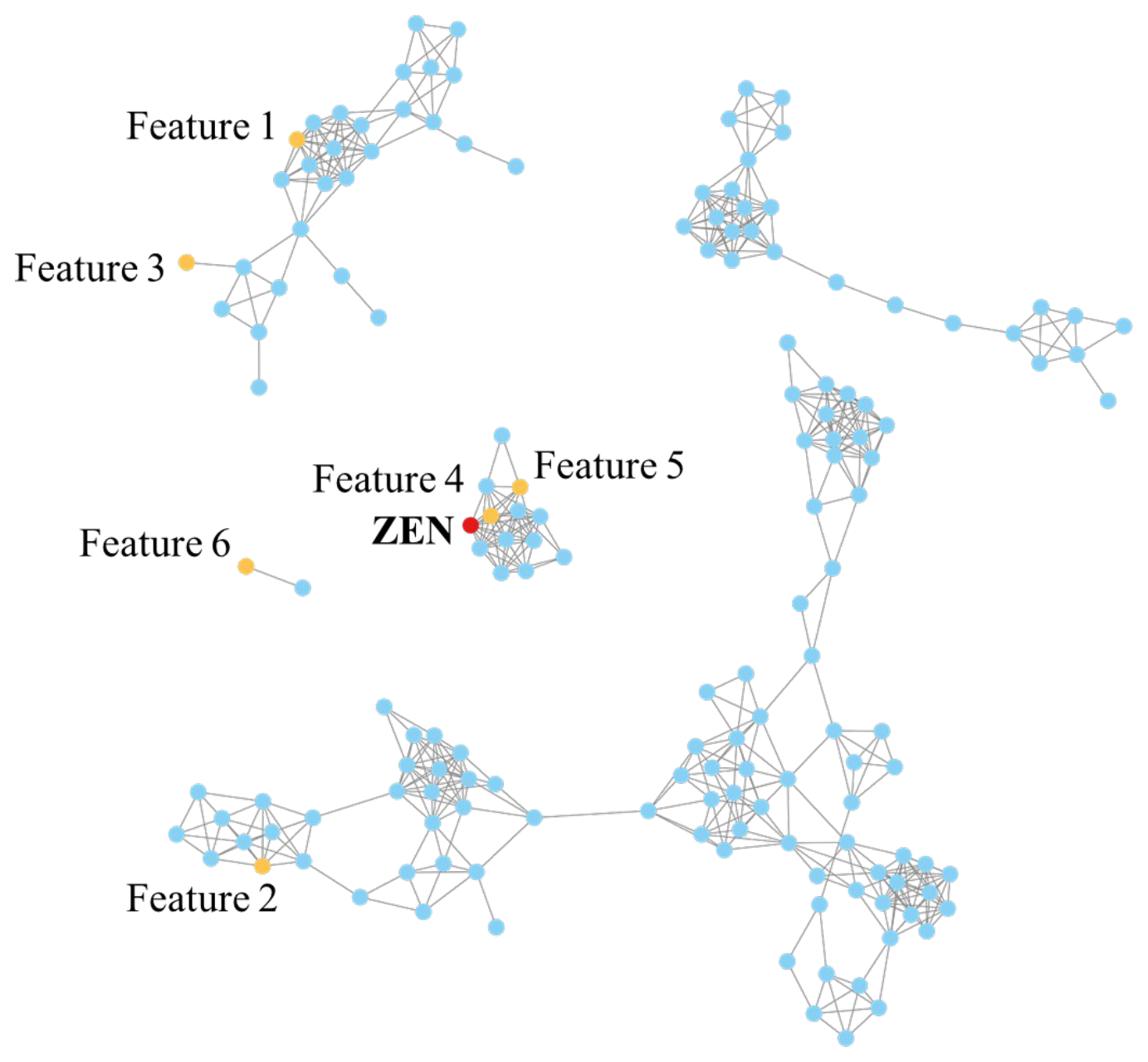

**Supplementary Fig. 4. Molecular-network context of zearalenone-related features.** Molecular-network components from the zearalenone (ZEN)-treated HepG2 spheroid conditioned-medium dataset. Nodes represent LC–MS/MS features, and edges represent spectral relationships meeting the modified cosine threshold of 0.70. Putative zearalenone glucuronide occurred in the ZEN-containing component, whereas putative zearalenol glucuronide occurred in a separate component. Both were retained by ANCHor through shared seed-anchor evidence. Red indicates ZEN, yellow indicates selected candidate features, and blue indicates other features. Feature numbers correspond to Supplementary Table 1.

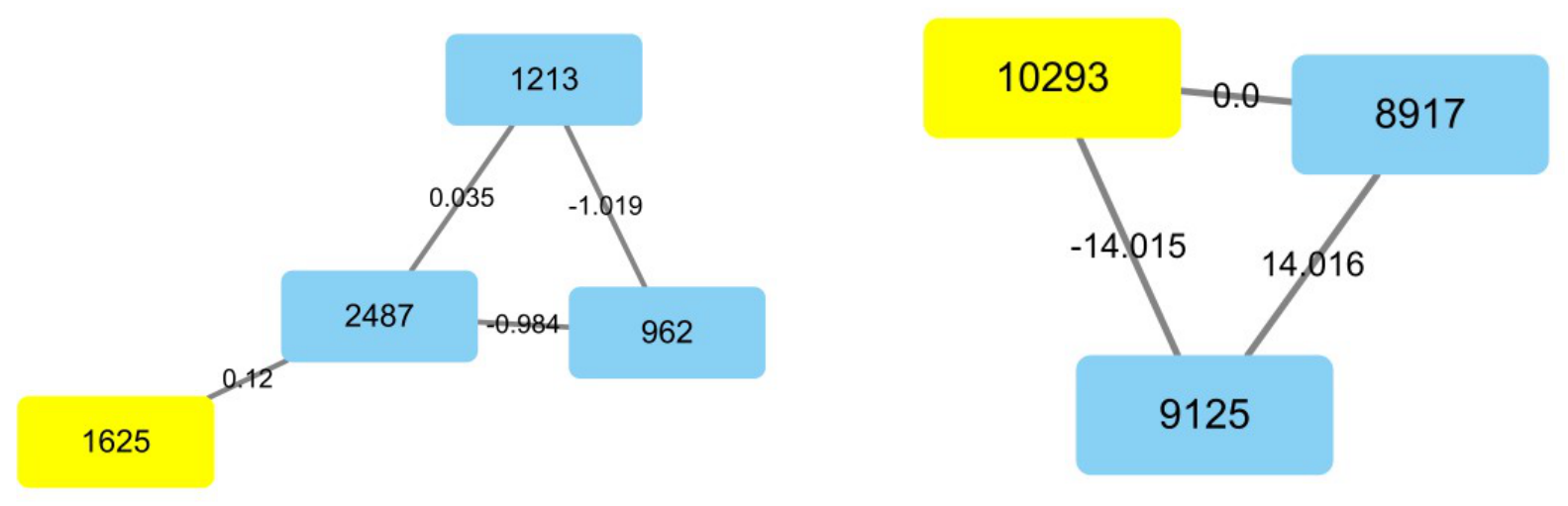

**Supplementary Fig. 5. Molecular-network context of two fecal unknown features.** Network components containing the seed at m/z 287.2439, subsequently identified as N¹,N¹²-diacetylspermine, and the seed at m/z 275.1025, investigated as a putative tryptophan–pyruvate-derived product. Yellow nodes indicate seeds, and blue nodes indicate other features. Node labels denote feature identifiers; edge labels indicate precursor m/z differences. Neither component contained a spectral-library-annotated neighboring feature.

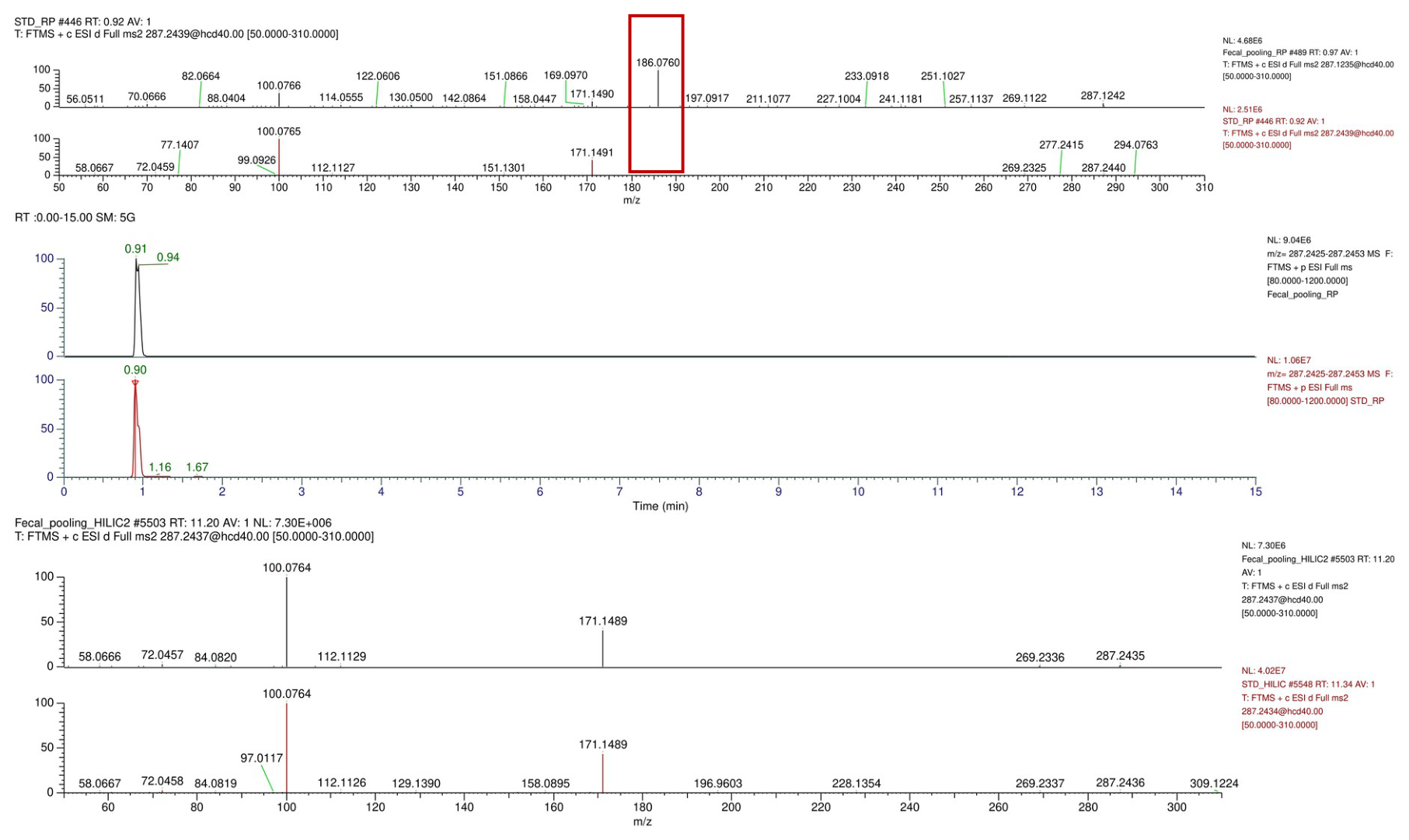

**Supplementary Fig. 6. Comparison with an authentic diacetylspermine standard.** Initial reversed-phase MS/MS comparison of the fecal feature at m/z 287.2439 and an authentic N¹,N¹²-diacetylspermine standard, extracted-ion chromatograms from separate reversed-phase injections, and MS/MS comparison after hydrophilic interaction liquid chromatography (HILIC) reanalysis. The additional fragment at m/z 186.076 in the initial fecal spectrum was attributed to co-isolation interference. HILIC reanalysis improved agreement between the feature and standard.

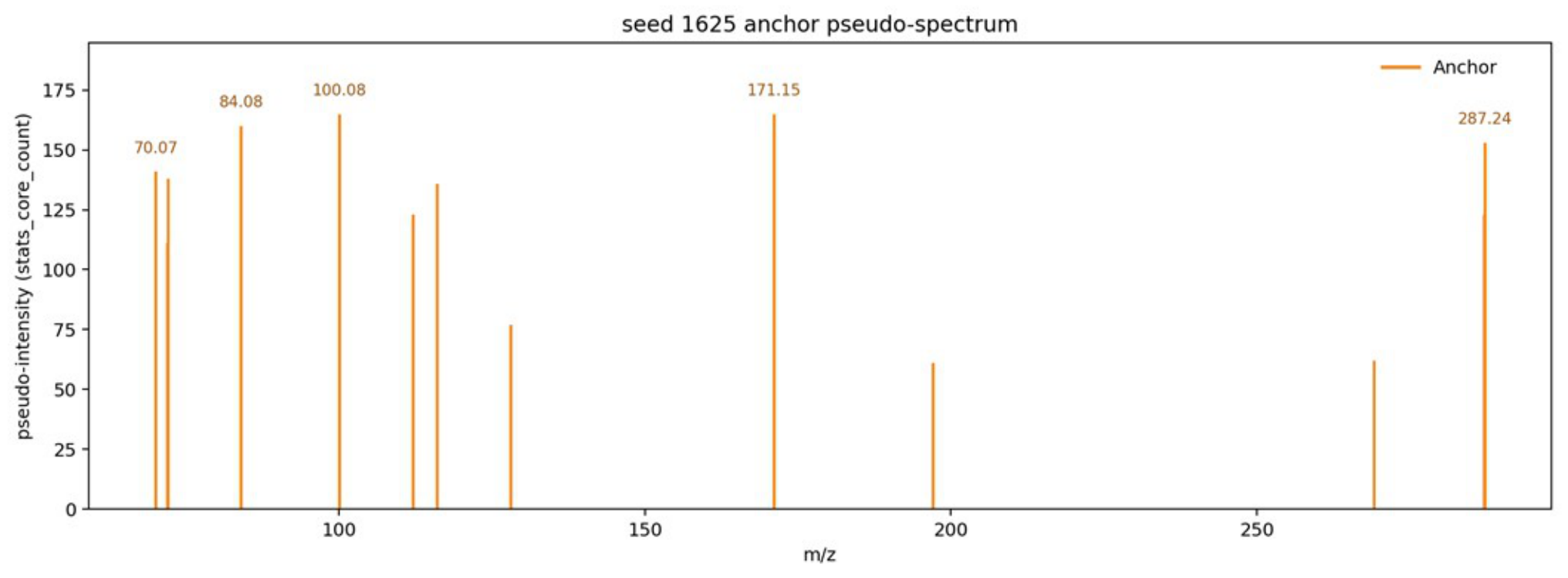

**Supplementary Fig. 7. Atlas-derived anchor evidence for the fecal polyamine seed.** Anchor pseudo-spectrum for the fecal seed at m/z 287.2439, subsequently identified as N¹,N¹²-diacetylspermine. Peak heights represent supporting atlas reference spectra rather than experimental signal intensity. The interference-associated fragment at m/z 186.076 in the initial fecal MS/MS spectrum was not selected as a final seed anchor.

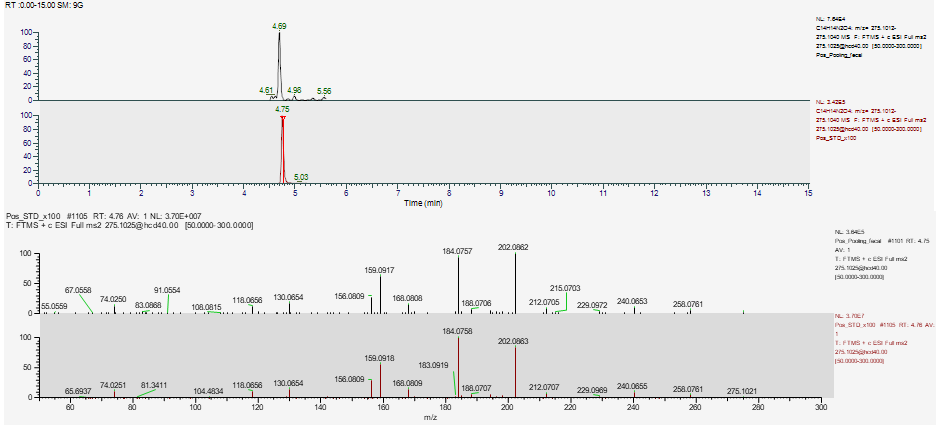

**Supplementary Fig. 8. Comparison with a tryptophan–pyruvate reaction product.** Extracted-ion chromatograms and MS/MS spectra of the fecal feature at m/z 275.1025 and a product generated by incubating L-tryptophan with pyruvic acid at 50 °C. Agreement in precursor m/z, retention behavior and fragmentation supported a tryptophan–pyruvate-derived tetrahydro-β-carboline annotation. These comparisons did not establish stereochemical configuration.

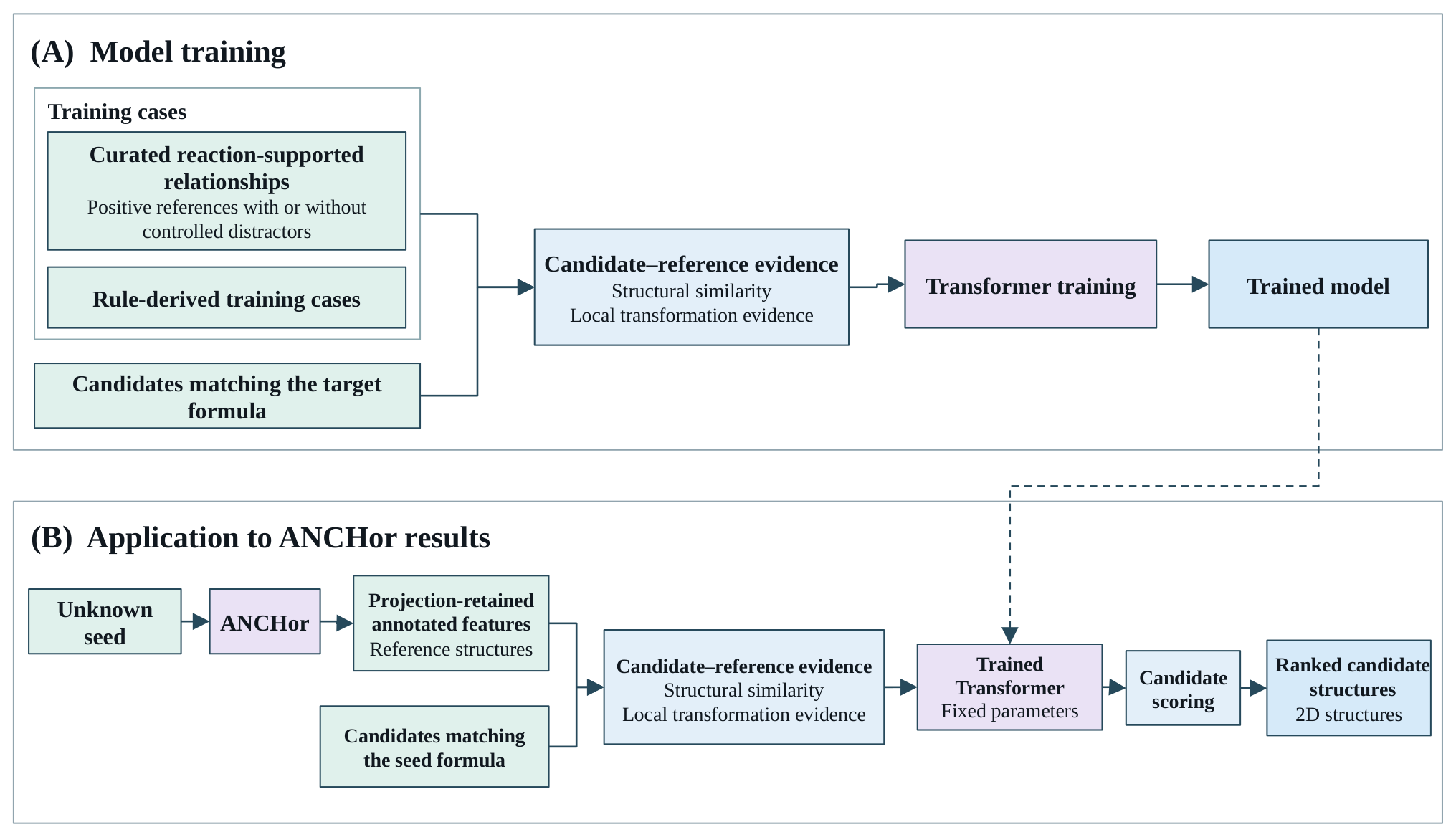

**Supplementary Fig. 9. Training and application of the optional prioritization model. a** Training used curated reaction-supported relationships, with or without controlled distractors, and additional rule-derived cases. Structural similarity and local transformation evidence for candidate–reference pairs trained the Transformer-based model to rank structures matching the target formula. Distractors represented incorrect proposed reference–target links. **b** During application, annotated features retained after ANCHor projection supplied reference structures. With parameters fixed, the model combined candidate–reference compatibility scores and relative reference weights to rank two-dimensional candidate structures matching the reconstructed seed formula.

### Supplementary Tables

Supplementary Table 1. Candidate ZEN biotransformation products.

| **Feature** | **Annotation** | **Modified cosine with ZEN** | **Phase** | **Transformation** | **Δm/z** | **Putative metabolite** |
| --- | --- | --- | --- | --- | --- | --- |
| Feature 1 | Zearalenol | 0.350 | I | hydrogenation(+H2) | 2.01565 | Zearalenol |
| Feature 2 | - | 0.452 | I + II | dehydrogenation(-H2) + methylation(+CH2) | 12 | - |
| Feature 3 | - | 0.396 | II | methylation(+CH2) | 14.01565 | - |
| Feature 4 | - | 0.896 | II | sulfation(+SO3) | 79.95682 | Zearalenone-4-sulfate |
| Feature 5 | - | 0.872 | II | glucuronidation(+C6H8O6) | 176.0321 | Zearalenone 14-glucuronide |
| Feature 6 | - | 0.090 | I + II | hydrogenation(+H2) + glucuronidation(+C6H8O6) | 178.0477 | zearalenol-14-glucuronide |

ZEN, zearalenone. Feature labels correspond to Supplementary Fig. 4. Proposed metabolite identities are putative.
